# Targeting STAT3 signalling using stabilised sulforaphane (SFX-01) inhibits endocrine resistant stem-like cells in ER-positive breast cancer

**DOI:** 10.1101/2020.02.03.932194

**Authors:** Bruno M. Simões, Angélica Santiago-Gómez, Chiara Chiodo, Tiago Moreira, Daniel Conole, Scott Lovell, Denis Alferez, Rachel Eyre, Katherine Spence, Aida Sarmiento-Castro, Bertram Kohler, Marilena Lanzino, Sebastiano Andò, Elisabetta Marangoni, Andrew H. Sims, Edward Tate, Sacha J. Howell, Robert B. Clarke

## Abstract

**PURPOSE:** Estrogen receptor (ER) positive breast cancer is frequently sensitive to endocrine therapy. Multiple mechanisms of endocrine therapy resistance have been identified, including cancer stem-like cell (CSC) activity. Sulforaphane (SFN) has previously been shown to target CSCs but its mechanism of action is unclear. Here we investigate SFX-01, a stabilised formulation of SFN, for its effects on breast CSC activity in ER+ preclinical models and to study its mechanism.

**EXPERIMENTAL DESIGN:** CSC activity was measured by mammosphere formation efficiency (MFE), aldehyde dehydrogenase (ALDH) activity, and tumor formation using patient samples and patient-derived xenograft (PDX) tumors treated with SFX-01 alone or in combination with tamoxifen or fulvestrant. Gene expression and SFN target proteins in treated samples were assessed.

**RESULTS:** SFX-01 reduced MFE of both ER+ primary and metastatic patient samples. Both tamoxifen and fulvestrant increased MFE and ALDH activity of PDX tumors, which was reversed by combination with SFX-01. SFX-01 significantly reduced tumor initiating cell frequency in secondary transplants at limiting dilution and reduced the formation of spontaneous lung micrometastases by PDX tumors in mice. Mechanistically, we establish that both tamoxifen and fulvestrant induce STAT3 phosphorylation. SFX-01 suppressed phospho-STAT3 and SFN directly bound STAT3 in patient and PDX samples. Analysis of ALDH+ cells from endocrine-resistant patient samples revealed activation of STAT3 target genes *MUC1* and *OSMR*, which were inhibited by SFX-01 in patient samples. Increased expression of these genes after 3 months’ endocrine treatment of ER+ patients (n=68) predicted poor prognosis.

**CONCLUSIONS:** Our data establish the importance of STAT3 signaling in CSC-mediated resistance to endocrine therapy and the potential of SFX-01 for improving clinical outcomes in ER+ breast cancer.

## INTRODUCTION

Three out of four cases of breast cancer express the estrogen receptor alpha (ER) and are treated with endocrine therapies, such as selective ER modulators (SERMs, e.g. tamoxifen), aromatase inhibitors (AIs, e.g. letrozole) and selective ER down-regulators (SERDs, e.g. fulvestrant) (1). However, despite the undoubted success of endocrine treatments, distant breast cancer recurrences and death occur at a steady rate for at least 15 years after the 5-10 year treatment period is completed (2). This finding stresses the need for new approaches that can provide long-term disease-free survival.

Endocrine resistance hampers the cure of ER+ breast cancer, and therefore major efforts have been employed to address the mechanisms. Acquired resistance can be mediated by modulation of ER activity through its mutation, by up-regulation of ER co-activators (e.g. FOXA1), by activation of mitogenic signalling pathways (e.g. MAPK, PI3K/AKT) induced by receptor tyrosine kinase activity (e.g. EGFR, ERBB2) or by overexpression of substrates for cyclin-dependent kinases (CDKs) (3, 4). Several drugs targeting these pathways have been tested in clinical studies, and recently CDK4/6 kinase inhibitors have been shown to increase overall survival (OS) of patients with advanced ER+ breast cancer and three (palbociclib, ribociclib and abemaciclib) are now FDA approved (5–7).

Cancer stem cells (CSCs) can be responsible for tumor initiation and growth and are more resistant than non-CSCs to cancer therapies, such as chemo- and radiotherapy (8). We and others have shown that breast CSCs (measured by the percentage of ALDH+, or mammosphere-forming cells) are not targeted by endocrine therapies in ER+ breast cancer (9–11). This leads to enrichment in cells with breast CSC activity which are dependent upon developmental signalling pathways, such as Notch and Wnt (12, 13). Eradication of endocrine-resistant CSCs is likely to provide long-term disease-free survival but so far none of the approved drugs for patients with ER+ tumors has been shown to target CSCs.

Sulforaphane (SFN), an isothiocyanate found in cruciferous vegetables, has demonstrated activity against breast CSCs (14), although clinical application has been hampered due to its inherent instability. In order to improve its stability, SFX-01, SFN conjugated within an alpha-cyclodextrin complex, has been developed (15).

Here, we report that SFX-01 used in combination with endocrine therapies prevents breast CSC enrichment in patient samples and PDX tumors in vivo, and mechanistically it directly targets STAT3 to inhibit its activity in endocrine resistance. STAT3 activation in patient primary tumors measured by expression of STAT3-induced genes predicts resistance to endocrine treatments.

## RESULTS

### SFX-01 reduces breast CSC activity in primary and metastatic ER+ breast cancer patient derived-samples

Mammosphere forming efficiency (MFE) and ALDH1 enzymatic activity have previously been shown to be a characteristic of breast CSCs (16, 17), which can be targeted by SFN in breast cancer cell lines (14). SFX-01 (SFN stabilised in alpha-cyclodextrin complex, **Figure 1A**) reduced MFE in thirteen of sixteen patient-derived ER+ tumor samples (**Figure 1B** and **Tables S1 and S2** for patient and tumor characteristics). Next, we assessed CSC activity following SFX-01 pre-treatment of cells from patients with endocrine-resistant breast cancer. SFX-01 reduced the percentage of ALDH+ cells in all six samples (and by >60% in five; **Figure 1C**) and also significantly decreased MFE in four of these samples (**Figure 1D**). These data suggest that short term SFX-01 treatment targets breast CSC activity in patient-derived and endocrine-resistant cells.

**Figure 1:**
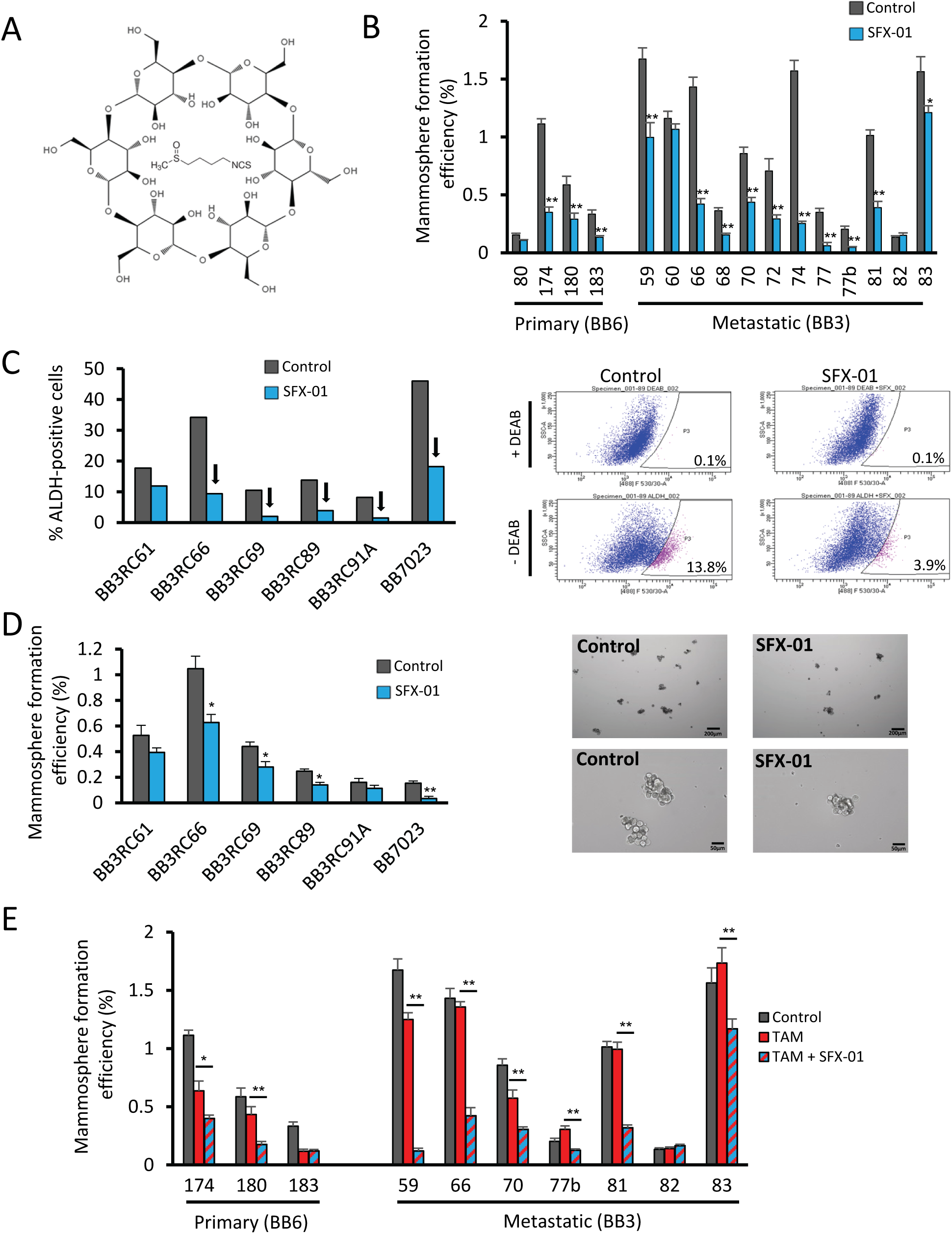
SFX-01 reduces breast CSC activity in primary and metastatic ER+ breast cancer patient derived-samples. A) Scheme of synthetic sulforaphane molecule stabilised in alpha-cyclodextrin complex as SFX-01. B) Mammosphere formation efficiency (MFE) of freshly isolated ER+ early (primary) and metastatic (pleural effusions and ascites) patient-derived samples cultured in the presence of SFX-01 (5 μM) or vehicle control. MFE data for each individual patient sample is represented. MFE was determined on day 7-9 and calculated by dividing the number of mammospheres formed (≥ 50μm diameter) by the original number of single cells seeded (500 cells/cm^2^) and is expressed as the mean percentage of mammosphere formation. C) Percentage of ALDH-positive cells in six ER+ metastatic BC patient-derived samples. Cells were grown in low-adherent plates in the presence of SFX-01 or water (control) for 72 hours and then subject to ALDEFLUOR assay. Arrows indicate a reduction greater than 60% compared to control. Representative FACS plots of ALDEFLUOR assay are shown. ALDH-positive cells were discriminated from ALDH-negative cells using the ALDH inhibitor, DEAB. D) Percentage of MFE in six ER+ metastatic BC patient-derived samples. Cells were pre-treated in low-adherent plates in the presence of SFX-01 or water (control) for 72 hours and then plated for MFE colony assay without drugs. Representative micrographs of mammospheres are shown. E) Percentage of MFE of ER+ early (primary) and metastatic patient-derived samples treated with tamoxifen alone (1 μM) or in combination with SFX-01 (5 μM). MFE data is represented as mean percentage ± SEM. * p < 0.05; ** p < 0.01

We then investigated the effect of SFX-01 on MFE of patient-derived ER+ tumor cells co-treated with tamoxifen. SFX-01 remained effective and significantly reduced MFE in combination with tamoxifen in eight out of ten samples tested (**Figure 1E**). In addition, we tested 3-day pre-treatment of ER+ cell lines (MCF-7, T47D and ZR-75-1) with tamoxifen or fulvestrant in combination with SFX-01. Both ALDH activity and MFE were reduced by SFX-01 in all cell lines confirming reversal of the breast CSC activity that is induced by short-term treatment with anti-estrogen drugs (**Supplementary Figure 1**).

### SFX-01 prevents tamoxifen enrichment for cells with cancer stem cell properties in patient-derived xenograft tumors

To better model the clinical scenario of our treatments, two ER+ patient-derived xenograft (PDX) models were grown subcutaneously in NSG mice, one from an early (HBCx34, (18)) and another from a metastatic tumor (BB3RC31, (19)). We assayed proliferation and breast CSC activity in these PDX models after 14-day *in vivo* treatment with either SFX-01, tamoxifen or combination of both drugs (**Figure 2A**). We observed that tamoxifen treatment decreases the proliferation marker Ki67 and tumor growth in both PDX models and SFX-01 reduced tumor growth in the HBCx34 model, but had no significant impact on BB3RC31 (**Figure 2B, Supplementary Figure 2A**). However, breast CSC activity measured by ALDH enzymatic activity or MFE was increased by tamoxifen and decreased by SFX-01, both alone and in combination with tamoxifen (**Figures 2C** and **2D**). We then performed *in vivo* limiting dilution transplantation of HBCx34 PDX cells derived from tumors previously treated with tamoxifen and/or SFX-01. Cells from tumors treated with SFX-01 could not form any tumors on reimplantation of 4,000 cells, in contrast with cells from tumors treated with vehicle control or tamoxifen alone (**Figure 2E**). Graphs representing tumor size observed in all serial dilutions are shown in **Supplementary Figure 2B**. Overall, tumors treated with SFX-01 had reduced tumor-initiating cell frequency confirming that SFX-01 targets breast CSC activity *in vivo* (**Figure 2E**).

**Figure 2:**
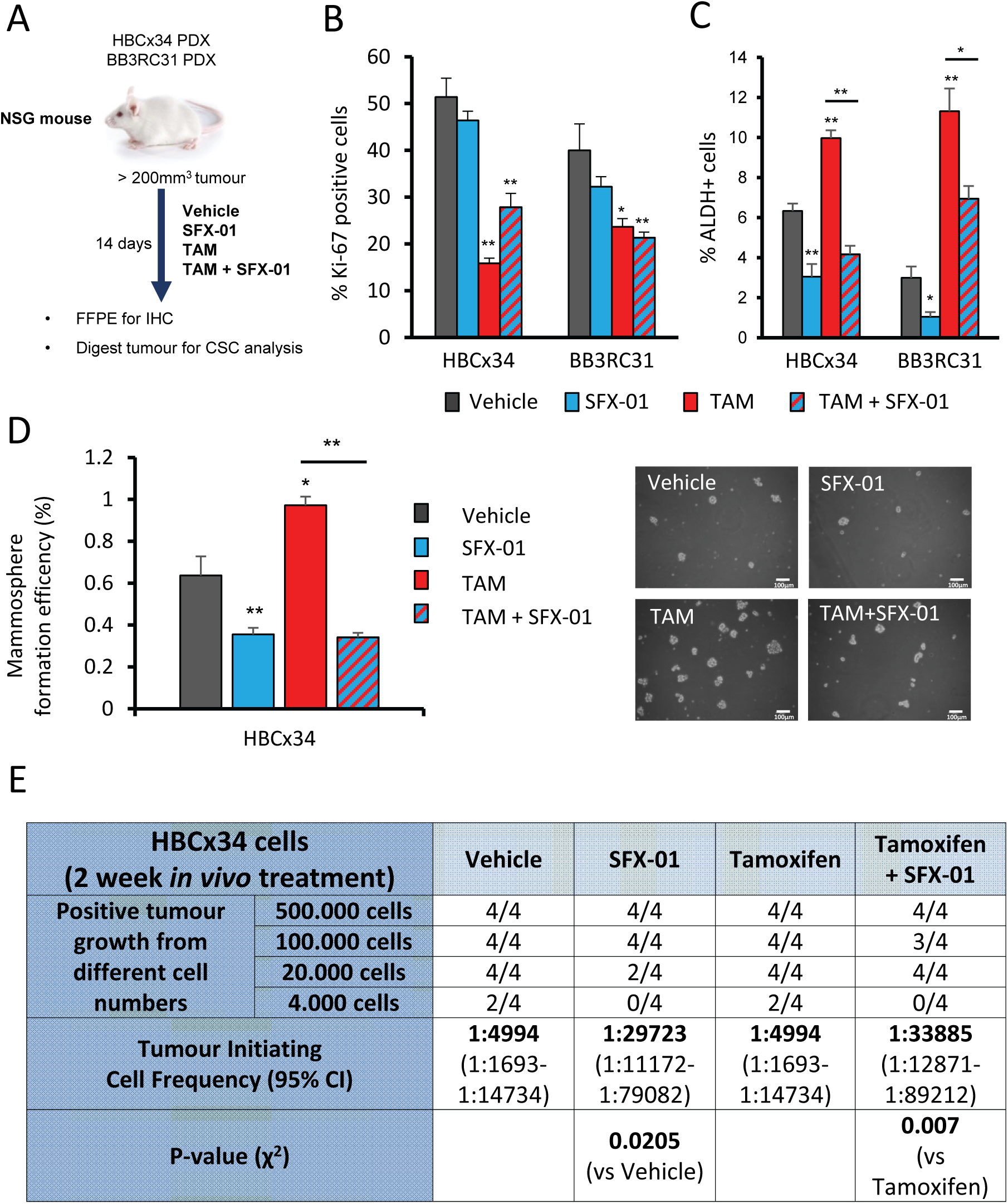
SFX-01 prevents tamoxifen enrichment for cells with cancer stem cell properties in patient-derived xenograft tumors. A) BB3RC31 and HBCx34 patient derived xenografts (PDXs) treated *in vivo* for 14 days with SFX-01 (300mg/kg/day, oral gavage) in the presence or absence of tamoxifen (10mg/kg/day, oral gavage). HBCx34 model was kindly provided by Dr Elisabetta Marangoni (Institute Curie, Paris). B) Quantification of Ki67 expression determined by immunohistochemistry showing that tamoxifen but not SFX-01 significantly decreases proliferation marker Ki67. C) Percentage of ALDH-positive cells was determined with ALDEFLUOR assay. ALDH-positive cells were discriminated from ALDH-negative cells using the ALDH inhibitor, DEAB. Mouse cells were excluded from the FACS analysis with anti-mouse MHC Class I (H-2K^d^) antibody. D) Mammosphere formation efficiency was determined on day 7-9 and calculated by dividing the number of mammospheres formed (≥ 50μm diameter) by the original number of single cells seeded (500 cells/cm^2^) and is expressed as the mean percentage of mammosphere formation. Representative micrographs are shown. E) Secondary transplantation of 500K, 100K, 20K and 4K cells after *in vivo* treatments. Experiment was carried out in NSG mice with 90-day slow-release estrogen pellets. Tumor growth (>75 mm^3^) was assessed at day 90 and is represented as mice positive for growth/mice tested for each cell number tested. ELDA of tumor-initiating cell frequency is shown. Data is represented as mean ± SEM. * p < 0.05; ** p < 0.01

### SFX-01 inhibits PDX tumor growth, prevents CSC activity and prevents spontaneous metastasis to the lungs

To further validate the *in vivo* effects of combining SFX-01 with anti-estrogens on CSC activity we treated HBCx34 PDX cells for 56 days with either tamoxifen or fulvestrant (**Figure 3A**). As expected, both anti-estrogens strongly reduced the number of proliferative cells measured by Ki67 expression (**Figure 3B**) whilst increasing both ALDH activity and MFE (**Figure 3C** and **3D**). SFX-01 had no effect on proliferation (**Figure 3B**) but significantly inhibited anti-estrogen stimulated breast CSC activity (**Figures 3C** and **3D**). 56-day treatment of the second PDX model (BB3RC31) also demonstrated that the anti-estrogen enrichment of breast CSCs can be reduced by SFX-01 (**Supplementary Figure 3A** and **3B**). Over the 8-week treatment period, tamoxifen plus SFX-01 significantly suppressed tumor growth versus tamoxifen alone in the HBCx34 (**Figure 3E**) but not the BB3RC31 PDX models (data not shown). On the other hand, fulvestrant treatment maintained tumor growth suppression at 8 weeks and no additive effect was seen with SFX-01 in either HBCx34 (**Figure 3F**) or BB3RC31 (data not shown) PDX models.

**Figure 3:**
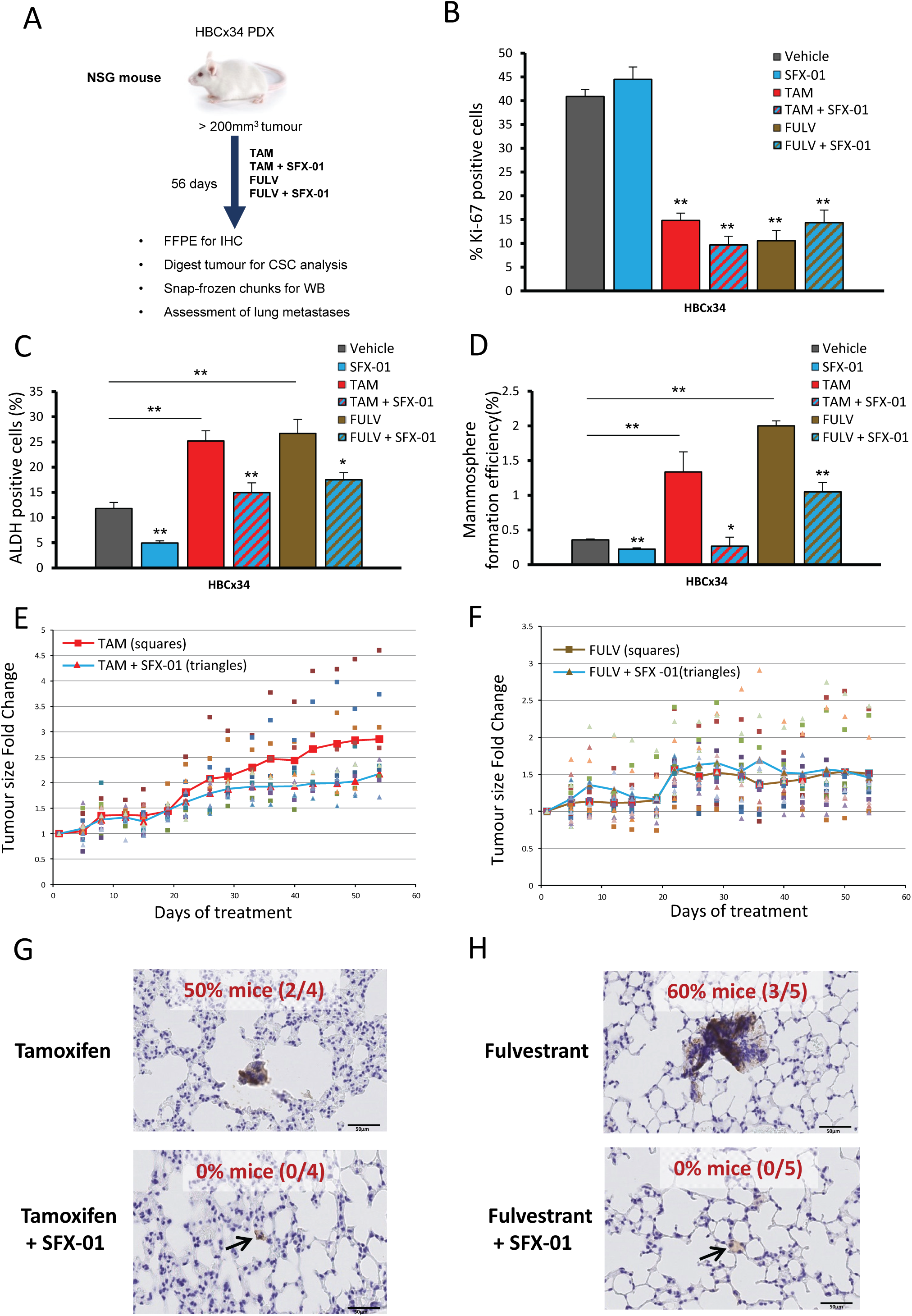
SFX-01 prevents CSC activity of PDX cells treated with tamoxifen and fulvestrant. SFX-01 inhibits tumor growth compared to tamoxifen alone and prevents formation of micrometastasis in the lungs. A) HBCx34 PDX treated *in vivo* for 56 days with tamoxifen (10mg/kg/day, oral gavage) or fulvestrant (200mg/kg/week, subcutaneous injection) in the presence or absence of SFX-01 (300mg/kg/day, oral gavage). B) Quantification of Ki67 expression determined by immunohistochemistry. C) Percentage of ALDH-positive cells was determined with ALDEFLUOR assay. ALDH-positive cells were discriminated from ALDH-negative cells using the ALDH inhibitor, DEAB. Mouse cells were excluded from the FACS analysis with anti-mouse MHC Class I (H-2K^d^) antibody. D) Mammosphere formation efficiency was determined on day 7-9 and calculated by dividing the number of mammospheres formed (≥ 50μm diameter) by the original number of single cells seeded (500 cells/cm^2^) and is expressed as the mean percentage of mammosphere formation. E-F) Tumor growth of HBCx34 PDX tumors treated *in vivo* for 56 days. Individual tumors (n=10) treated with tamoxifen (E) or fulvestrant (F) are represented by squares and tumors treated in combination with SFX-01 are represented by triangles. Red line shows average tumor growth for tamoxifen (E) or fulvestrant (F) and blue line shows average tumor growth for the combination treatment with SFX-01. G-H) Mice lungs were stained with anti-human mitochondrial antibody and micrometastases with at least 10 cells were counted. Percentage of mice bearing micrometastases for each tamoxifen (G) and fulvestrant (H) treatment group is shown. Data is represented as mean ± SEM. * p < 0.05; ** p < 0.01

We next investigated whether treatments had an effect on PDX spontaneous lung metastasis. Using a human-specific mitochondrial antibody we could clearly observe micrometastases in the lungs from mice bearing HBCx34 PDX tumors treated with tamoxifen or fulvestrant (**Figure 3G** and **3H**). In contrast, lungs from HBCx34 mice treated with tamoxifen or fulvestrant in combination with SFX-01 were free from micrometastases, although disseminated tumor cells persisted (**Figure 3G** and **3H**). In the BB3RC31 PDX model SFX-01 reduced micrometastases by 50% in combination with fulvestrant but no effect was seen in combination with tamoxifen (**Supplementary Figure 3C** and **3D**), suggesting SFX-01 hinders colonisation of the lungs by disseminated tumor cells in sensitive tumors.

### SFX-01 targets STAT3 signalling, which is activated by anti-estrogen therapy

We next sought to determine the mechanism of sensitivity to SFX-01. The direct binding targets of SFN have been assessed in breast cancer cell lines, identifying STAT3 and NF-κB subunits among the top high-affinity targets (20). STAT3 was a particularly interesting target since ALDH+ breast CSCs and tamoxifen resistant cells have previously been shown to express higher levels of phospho-STAT3 (active form), and inhibition of STAT3 reduces ALDH+ cell numbers and tumorigenicity (21, 22). HBCx34 PDX tumors treated *in vivo* for 56 days with tamoxifen or fulvestrant showed increased phospho-STAT3 expression which was inhibited by co-treatment with SFX-01 (**Figure 4A**). In the BB3RC31 PDX model we also observed that fulvestrant, but not tamoxifen, increased phospho-STAT3 expression and SFX-01 only reduced phospho-STAT3 levels in fulvestrant-treated tumors (**Supplementary Figure 4B**). We also assessed whether phospho-NF-κB p65 was modulated in a similar way to STAT3 but overall its expression was not changed in either HBCx34 or BB3RC31 PDX treated tumors (**Supplementary Figure 4A** and **4B**). Next, we examined STAT3 activation in a HBCx34 PDX model that has been selected for tamoxifen resistance, which we previously demonstrated to display an enrichment of breast CSCs (12). Increased phospho-STAT3 was seen compared to the parental endocrine-sensitive HBCx34 PDX model and this was reversed by treatment with SFX-01 (**Figure 4B**). Pull-down experiments with biotin-labelled SFN in HBCx34 PDX tumor cells confirmed direct interaction of SFN with STAT3 in both anti-estrogen and vehicle-treated tumors (**Figure 4C**), establishing STAT3 as the likely target of SFX-01.

**Figure 4:**
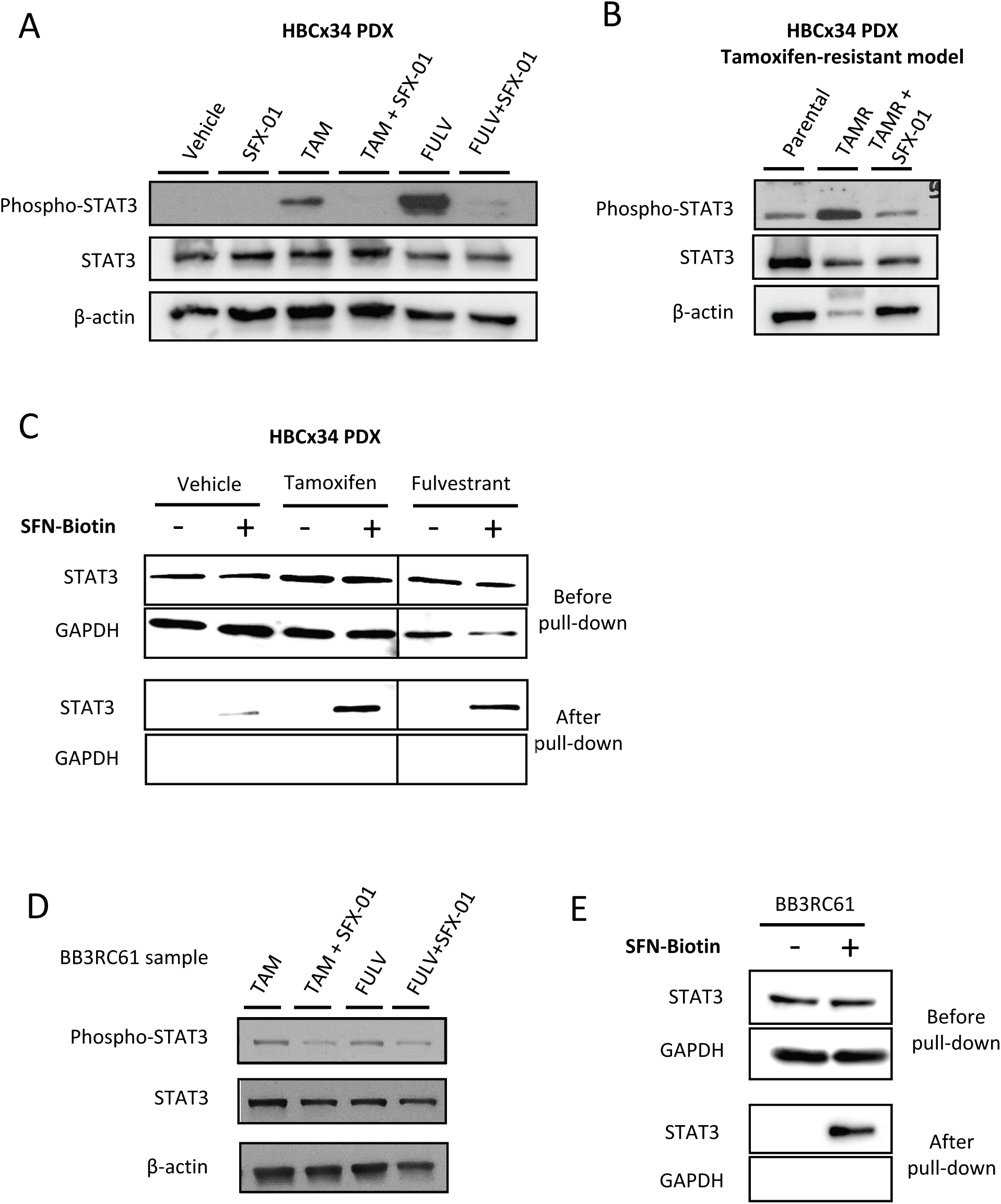
SFX-01 targets STAT3 signalling, which is activated by anti-estrogen therapy. A) phospho-STAT3 and total STAT3 protein expression levels determined by Western Blot in HBCx34 PDX treated *in vivo* for 56 days with tamoxifen or fulvestrant in the presence or absence of SFX-01. β-actin was used as a reference for the loading control. B) Protein expression levels in tamoxifen-resistant (TAMR) HBCx34 PDX treated *in vivo* for 56 days with SFX-01. C) Total STAT3 protein detection before and after SFN-Biotin-Avidin pull-down experiments in HBCx34 PDX treated tumors. GAPDH was used as a reference for the loading control. D) Protein expression levels in metastatic patient-derived sample BB3RC61 treated for 72Hours with tamoxifen or fulvestrant in the presence or absence of SFX-01. E) Total STAT3 protein detection before and after SFN-Biotin-Avidin pull-down experiments in BB3RC61 sample. GAPDH was used as a reference for the loading control.

Next we tested whether SFX-01 inhibited STAT3 signalling in anti-estrogen resistant patient samples. Cells from metastatic sample BB3RC61 treated with SFX-01 showed a significant reduction in phospho-STAT3 expression compared to tamoxifen or fulvestrant treatments alone (**Figure 4D**). Furthermore, we observed direct binding of SFN to STAT3 in cells from this same patient sample (**Figure 4E**). Analysis of four additional anti-estrogen resistant metastatic samples confirmed that SFX-01 combined with either tamoxifen or fulvestrant inhibited STAT3 activity in three of these patient samples (BB3RC44, BB3RC66 and BB7103), though SFX-01 only inhibited fulvestrant-induced STAT3 activation in BB3RC66 sample (**Supplementary Figure 4C**). Interestingly, SFX-01 did not repress STAT3 activity in sample BB7121 that had previously been treated clinically with SFX-01 (STEM trial). This patient’s disease had progressed in the presence of SFX-01 and was thus resistant to it.

On the whole, we have established, using PDX models *in vivo* and patient samples *in vitro*, that SFX-01 targets STAT3 and can inhibit its activity that is induced by treatment with anti-estrogens.

### STAT3-related genes are up-regulated in anti-estrogen resistant ALDH+ cells and are associated with worse outcomes for ER+ breast cancer patients

To investigate which genes are regulated by STAT3 in the anti-estrogen resistant ALDH+ CSC population, we compared the gene expression profile between ALDH+ and ALDH-cells of 4 metastatic ER+ patient samples treated with endocrine therapy prior to analysis (**Supplementary Figure 5A**). Ingenuity Pathway Analysis (IPA) demonstrated differential expression (fold change ≥ 2) of 28 STAT3-related genes that were all up-regulated in ALDH+ cells (**Figure 5A**). Next, we hypothesised that this 28-gene STAT3 signature of ALDH+ cells could predict prognosis of patients diagnosed with ER+ breast cancers. In published gene expression microarray datasets from 762 ER+ tumors (KMplotter, (23)), we found that elevated expression of the 28 STAT3-related genes before treatment was significantly associated with breast cancer recurrence (**Figure 5B**). Thus, activation of STAT3 signalling in ER+ tumors is associated with more aggressive disease.

**Figure 5:**
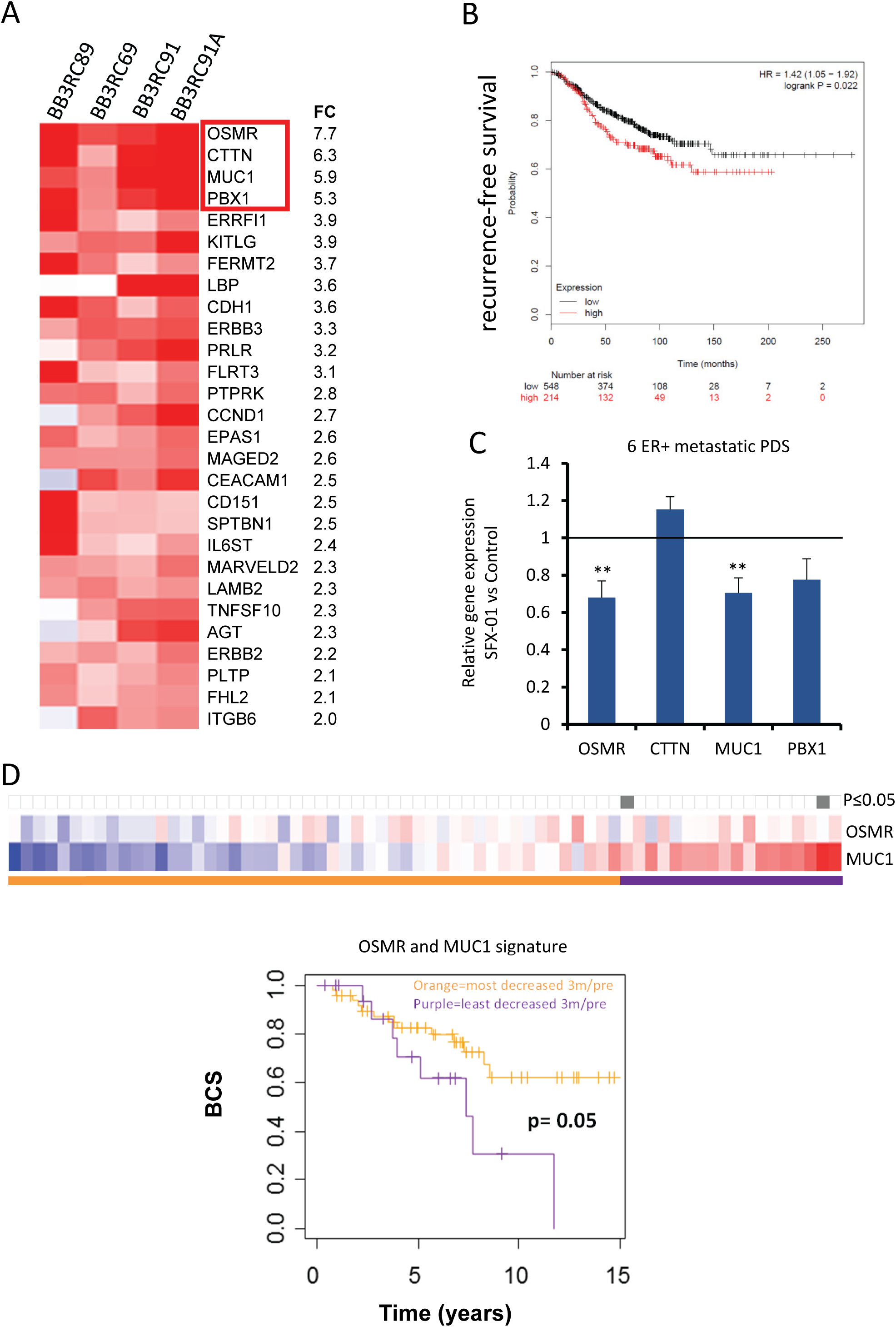
STAT3-related genes are up-regulated in anti-estrogen resistant ALDH+ cells and are associated with worse outcomes for ER+ breast cancer patients. A) Heatmap illustrating the 28 STAT3-related genes differentially expressed in ALDH+ cells of four ER+ metastatic patient derived cells treated with anti-estrogen treatments (fold change ≥ 2, pairwise Rank Products). Red colour represents gene up-regulation while blue shows down-regulation in ALDH+ relative to ALDH-cells. B) Kaplan-Meier analysis of high versus low expression of STAT3 28-gene signature showing recurrence-free survival in ER+ breast cancer patients. The gene expression data is from published microarray datasets available in KMplotter (23). C) Expression of *OSMR, CTTN, MUC1* and *PBX1* genes was assessed by real-time qPCR analysis and compared to control to determine fold change. Metastatic patient-derived cells were treated for 72 hours with SFX-01 or water (control). D) Heatmap showing *OSMR* and *MUC1* gene expression in patient matched dataset after 3 months treatment with letrozole relative to baseline. Heatmap is ranked from left to right using the sum of the expression of the two genes. Red indicates up-regulation and blue indicates down-regulation in treated tumors relative to baseline. All significant cut-points (p ≤ 0.05) are shown in grey. The Kaplan-Meier plot demonstrates that elevated expression of these genes is significantly associated with decreased breast cancer specific survival (BCS). Vertical bars on survival curves indicate censored cases. p value is based on a log-rank (Mantel-Cox) test.

We therefore tested whether SFX-01 could reduce expression of the top four STAT3 related genes in ER+ anti-estrogen resistant patient samples, and we found that two of the genes (*OSMR* and *MUC1*) were significantly down-regulated by SFX-01 treatment (**Figure 5C**). Based on these findings, we investigated if failure to reduce expression of these two STAT3 genes with anti-estrogen treatments was associated with worse outcomes due to STAT3 activation in a 68-patient matched dataset (24) before and after 3 months treatment with an aromatase inhibitor. Remarkably, increase in the sum of these two genes (*OSMR* and *MUC1*) upon anti-estrogen treatment was significantly associated with decreased breast cancer specific survival (**Figure 5D**). The change in expression of each of these genes individually was not significantly associated with outcome (data not shown).

Thus, persistent activation of the STAT3 signalling pathway after anti-estrogen treatment could potentially be used as a biomarker to predict endocrine resistance and, hence, to identify the patient group most likely to benefit from STAT3 inhibition with SFX-01 therapy.

## DISCUSSION

In this study we establish that SFX-01 targets anti-estrogen resistant breast CSCs in patient samples and patient-derived xenografts (PDXs). Compared with anti-estrogen treatment alone, SFX-01 combined with anti-estrogen drugs significantly reduced ALDH+ CSCs, their sphere-forming activity and tumor initiating cell frequency, as well as formation of lung micrometastases in PDX tumors grown in mice. Mechanistically, both tamoxifen and fulvestrant induced STAT3 phosphorylation although not always to the same degree in individual tumors. We show that SFX-01 suppresses this adaptive STAT3 phosphorylation and establish that SFN directly binds to and targets STAT3 in cells from patient samples and PDX tumors. Importantly, analysis of ALDH+ cells from endocrine-resistant patient samples revealed activation of STAT3 signalling and expression of STAT3 target genes *MUC1* and *OSMR*, which were down-regulated by SFX-01 in patient samples. Moreover, increased expression of these two genes after 3 months’ endocrine treatment predicted poor prognosis in patients with ER+ tumors. We propose that inhibiting STAT3 signalling with SFX-01 may help to overcome endocrine therapy resistance and recurrence in ER+ breast cancer.

We previously reported that CSC activity is increased by the anti-estrogens tamoxifen and fulvestrant, both *in vitro* and *in vivo* (12), and others have demonstrated a role for SFN in targeting breast CSCs. Here, we show a stabilised form of SFN, SFX-01, is a potent inhibitor of endocrine resistant CSCs. In contradistinction to the previous report that used cell lines (14), we establish that STAT3, not Wnt signalling, is the primary target mechanistically responsible for endocrine resistance of breast CSCs in human breast tumors.

STAT3 activation has been associated with ALDH expression in triple negative breast cancer cell lines (22), however, this is the first time this association has been reported for endocrine resistant breast cancer. We previously reported that Notch4 signalling is increased in anti-estrogen resistant ALDH+ breast CSCs (12), and using a tamoxifen-resistant MCF7 sub-line, an association was reported between Notch4 and STAT3 activation (21). However, we have not confirmed here that STAT3 phosphorylation is confined to the ALDH+ population. The relevance of our finding is strengthened by demonstrating that STAT3 is activated in both anti-estrogen treated PDX and patient-derived samples.

The recent finding that SFN bound directly to STAT3 with high-affinity in breast cancer cell lines (20) led us to investigate STAT3 as the target in anti-estrogen treated PDX and endocrine resistant patient-derived samples. We show convincingly that binding occurs and that SFX-01 reduces the phosphorylation of STAT3 *in vitro* and *in vivo*, in short-term anti-estrogen treatment and established resistance. This establishes its utility and provided a rationale for its use in clinical trials in patients. Therefore, we recently recruited 47 patients with metastatic breast cancer who were progressing on endocrine therapy and showed that the addition of SFX-01 to the endocrine therapy resulted in clinical benefit in 25% of patients (25). Our *in vivo* data demonstrate where primary tumor phospho-STAT3 is reduced, there is a reduction in metastatic lung colonisation. This indicates that SFX-01 could reduce distant recurrence rates in early breast cancer.

The *in vivo* data presented, demonstrating reduction in metastatic lung colonisation when primary tumors phospho-STAT3 was reduced, suggest that SFX-01 could reduce distant recurrence rates in early breast cancer. Although this clearly requires additional preclinical and clinical testing.

In order to better target STAT3-driven endocrine resistant breast cancer, we analysed gene expression in samples from advanced breast cancer patient who had received prior endocrine therapy. We derived a STAT3 signature consisting of 28 genes that was associated with poor progression-free survival of ER+ breast cancer patients. Importantly, two of the most expressed genes were down-regulated by SFX-01 treatment of patient-derived samples. These two genes, *OSMR* and *MUC1*, were both STAT3-responsive and SFX-01-sensitive in patient samples, and their increased expression after 3 months’ endocrine treatment accurately predicted survival over a 14 year period after surgery for breast cancer. These data suggest that this two gene signature could have potential to select tumors responsive to SFX-01, which would have to be confirmed in a larger group of patients.

*OSMR* has already been associated with worst prognosis although not specifically for ER+ tumors (26), and an association between OSMR signalling and reduced ER levels has been reported (26). In addition, there are reports showing that OSMR signalling increases invasive capacity of breast cancer cell lines, perhaps explaining increased metastasis (27). Moreover, a MUC1 signature is associated with high MUC1 expression and predicts failure to tamoxifen treatment (28).

Thus, overall we establish the potential of SFX-01 for clinically meaningful improvements to endocrine therapy in ER+ BC by inhibiting STAT3 signalling and reversing CSC-mediated resistance. The derivation of a STAT3 pathway signature that predicts poor prognosis and resistance to endocrine therapy indicates that a sub-population of breast tumors might be the rational targets for SFX-01 therapy in combination with current endocrine therapies.

## EXPERIMENTAL PROCEDURES

### Breast cancer patient-derived samples

Early breast cancer (BC) samples were collected from patients undergoing surgical resection of the primary tumor at three NHS Foundation Trusts (South Manchester, Salford Royal and The Pennine Acute Hospitals). Metastatic fluids (ascites or pleural effusions) drained from advanced breast cancer patients were collected at the Christie and South Manchester Hospitals NHS Foundation Trusts through the Manchester Cancer Research Centre Biobank. Fully informed consent from all patients was obtained in accordance with local National Research Ethics Service guidelines (study numbers: 05/Q1402/25, 05/Q1403/159 and 12/ROCL/01). Early and metastatic BC samples were processed as described previously (12, 19). Supplementary **Tables S1** (early BC) and **S2** (metastatic BC) present the clinico-pathological characteristics of the samples used in this study.

### *In vivo* experiments using patient-derived xenografts

All *in vivo* studies were carried out in accordance with the UK Home Office (Scientific Procedures) Act 1986. NOD *scid gamma* (NOD.*Cg-Prkdc*^scid^ Il2rg^tm1Wjl^/SzJ, NSG) and athymic Nude mice (Foxn1^nu^) mice (Charles River) were used in the following experiments.

Small fragments of HBCx34 (early) and BB3RC31 (metastatic) breast cancer patient-derived xenograft (PDX) tumors were implanted subcutaneously into the flanks of 8-12-week-old female NSG mice. These two pre-clinical models are estrogen-dependent, so animals were administered with 8 μg/ml 17-beta estradiol (Sigma-Aldrich, #E2758) in drinking water at least 4 days previously to implantation until the end of the experiment. HBCx34 PDX model and their endocrine resistant variant (tamoxifen-resistant, TAMR) were kindly provided by Dr Elisabetta Marangoni (Institut Curie, Paris) (18). *In vivo* experiments with HBCx34 TAMR PDX model were carried out using Nude mice in the Institut Curie. For clinical details of BB3RC31 refer to (12, 19).

PDX tumors were treated when reached 200-300 mm^3^. Tumor size and animal weight were recorded twice weekly. SFX-01 (Evgen Pharma, 300 mg/kg/day) and tamoxifen citrate (Sigma-Aldrich, #T9262, 10 mg/kg/day) were administered by oral gavage (0.1 ml/dose) on a five day from seven day basis (weekends excluded) for 14 or 56 days; whereas Fulvestrant (200mg/kg/week, Astrazeneca) was injected subcutaneously (0.1 ml/dose) on a weekly basis for 56 days. SFX-01 and tamoxifen citrate were made in 1% carboxymethylcellulose (Sigma-Aldrich, #C9481) dissolved in distilled water. Upon termination, xenografts tumors were collected in ice-cold DMEM media and processed as described in (12) for further downstream analyses. Mouse lungs were formalin-fixed paraffin-embedded for histological assessment of metastatic disease.

*In vivo* limiting dilution assays were performed to evaluate tumor initiation ability of HBCx34 PDX after two weeks of *in vivo* treatment. Xenografts treated with either SFX-01, tamoxifen citrate, combination or vehicle control for 14 days were collected and digested using collagenase-hyaluronidase (Stem Cell Technologies) to obtain single-cell suspensions. Serial limiting dilution of PDX-derived cells (500,000; 100,000; 20,000; 4,000 cells) were resuspended in mammosphere media:Matrigel (1:1) and subcutaneously injected into the flank of NSG mice (n=4 per condition). 90-day slow-release estrogen pellets were implanted subcutaneously two days prior to cell injection (0.72 mg, Innovative Research of America). At day 90 after cell injection, positive tumor growth was considered in mice bearing a tumor greater than 75 mm^3^. The tumor-initiating cell frequency was calculated using Extreme Limiting Dilution Analysis (ELDA) software (The Walter and Eliza Hall Institute of Medical Research) with a 95 % confidence interval. p-values were obtained by Chi-squared statistical analysis.

### Cell lines and reagents

MCF7, T47D and ZR-75-1 cell lines were obtained through the American Tissue Culture Collection (ATCC) and cultured in DMEM/F12, GlutaMAX (Gibco) containing 10% foetal bovine serum (Gibco). Cells were pre-treated in adherence for 3 days with either ethanol (vehicle control), 1μM 4-OH-tamoxifen (Sigma-Aldrich) or 0.1μM fulvestrant (TOCRIS Bioscience) in the presence of 5μM SFX-01 (Evgen Pharma) or water (vehicle control), respectively.

### Mammosphere colony assay

Cancer stem cell activity using the mammosphere colony assay was determined following the protocol described elsewhere (29). Freshly isolated cells from ER+ primary and metastatic patient-derived samples (500 cells/cm^2^) were cultured for 7-9 days in mammosphere culture conditions. When indicated, cells were either treated directly in the mammosphere media or pre-treated for 72h in low adherence previously to assess their mammosphere forming ability. PDX-derived cells (500 cells/cm^2^) were cultured for 7-10 days. Cell lines were pre-treated in adherence for 3 days and then cultured in suspension for 5 days (200 cells/cm^2^). Mammosphere formation efficiency (MFE, %) was calculated by dividing the number of mammospheres formed per well (≥ 50μm diameter) by the number of single cells seeded per well. MFE is expressed as the average percentage of MFE.

### ALDEFLUOR Assay

Cancer stem cell activity was measured using the ALDEFLUOR assay (Stem Cell Technologies). This fluorescent reagent system allows detecting the activity of aldehyde dehydrogenase (ALDH), enzyme highly active in mammary stem cells (16). The assay was performed following the manufacturer’s instructions. Single cells were resuspended in ALDEFLUOR assay buffer containing 1.5 mM bodipyaminoacetaldehyde (BAAA), ALDH substrate, and incubated in the dark for 45 min at 37 °C. In parallel, a fraction of the cells was incubated in the presence of a 2-fold molar excess of the ALDH inhibitor, diethylaminobenzaldehyde (DEAB), in order to correct for background staining and define the ALDH positive region during the analysis. When using PDX tumors, a Pacific Blue anti-mouse H-2K^d^ MHC Class I antibody (BioLegend, #116616) was used to remove mouse cells from the analysis. In all cases, dead cells were excluded using 7-aminoactinomycin D (7AAD, BD Biosciences). Data were acquired with the BD LSRII flow cytometer (BD Biosciences) and analysed using the BD FACSDiva^TM^ software.

### Western blot

Cells were resuspended in Protein Lysis Buffer (25 mM HEPES, 50 mM NaCl, 50 mM sodium pyrophosphate, 50 mM sodium fluoride, 1% Triton-X-100, 10% glycerol, 5 mM EDTA, Roche MiniProtease Inhibitor cocktail tablet, 1 μM PMSF) and placed on a rotator for 1 hour at 4°C. Protein lysates were then obtained by centrifugation at 10,000×g for 10 min at 4°C. Protein concentration was determined using the BCA Protein Assay kit (Thermo Fisher Scientific). Samples were loaded in pre-cast 10% gels for SDS-PAGE (BioRad, #456-1033) and run at 200 V for 1 hour. Proteins were then wet-transferred to a 0.2 μm Nitrocellulose membrane (Amersham GE Healthcare) at 150 V for 1 hour. Membranes were blocked in 5% skimmed milk or bovine serum albumin (BSA)/TBS-0.001 % Tween 20 (TBST-T) for 1 hour at room temperature followed by an overnight primary antibody incubation at 4°C. Primary antibodies used were: Phospho-STAT3 Y705 (Cell Signalling, #D3A7), STAT3 (Cell Signalling, #124H6), phospho-NFκB p65 S536 (Cell Signalling, #93H1), NFκB (Cell Signalling, #D14E12), β-actin (Sigma, #A2228). Following 3 washes with TBS-T, membranes were incubated with HRP-conjugated secondary antibodies (Dako #P0447, #P0448, #P0449) for 1 hour at room temperature. Blots were developed with Luminata Classico or Luminata Forte Western HRP Substrate (EMD Millipore, #WBLUC0100, # WBLUF0100) by exposing the membranes to hyperfilm (Amersham GE Healthcare).

### Immunohistochemistry

Formalin-fixed paraffin-embedded PDX tumors and mouse lung tissue were cut into 4 μm-thick sections for immunohistochemical analysis. Using the Bond Automated Stainer (Leica Biosystems), tumor sections were stained for human Ki67 (Dako, #M7240) using the Target Retrieval Solution pH 9.0 (Dako, #S2367). The antibody was detected using Dako EnVision Detection System Peroxidase/DAB (Dako, #K5007). Definiens Tissue Studio software was used to quantify the staining. The percentage of positive epithelial cells was scored at least on 3 different tissue samples.

PDX lungs were examined for metastatic lesions of human origin by staining with anti-human mitochondrial antibody (Abcam, #ab92824). At least 2× step sections per lung were stained on a Bond Automated Stainer (Leica Biosystems) using standard protocol F and Target Retrieval Solution pH 6.0 (Dako) for 20 min. Direct visual examination of stained sections was carried out by an unbiased observer to detect micrometastases. Lesions with at least 10 cells were classed as micrometastases.

All sections were counterstained with haematoxylin.

### Affinity pull-down assays

Sulforaphane activity-based probe 2 (SFN-ABP) was prepared and used in affinity pull-down assays following the approach described in (20). Cells from patient-derived samples and xenografts were resuspended in serum-free DMEM media containing 5 μM SFN ABP probe and incubated for 30 min at 37°C. Following in-cell labelling with the probe, cells were washed with serum-free DMEM media and lysed in PBS pH 7.4 containing 1% Triton-X-100, 0.1% SDS, 1x Complete EDTA-free protease inhibitor tablet. After 5 min on ice, samples were sonicated and lysates were obtained upon centrifugation at 10,000 x g for 5 min at 4°C. Protein concentration was determined in the lysates and aliquots at 1 mg/ml were made with lysis buffer (100 μl per aliquot). Probe-labelled proteins were ligated to Copper-catalysed azide-alkyne cycloaddition (CuAAC), which was subsequently captured with azido-TAMRA-biotin (AzTB) (click reaction). Samples were vortexed for 1 hour at room temperature and then the click reaction was quenched by addition of 10 mM EDTA. Proteins were precipitated by centrifugation at 6,000×g for 4 min in methanol:chloroform:water (4:1:2). Protein pellets were then washed twice with 4 volumes methanol by vortexing, sonication and centrifugation at 8,000×g for 4 min. Protein pellets were air dried for 10 min before being resuspended in 0.2% SDS/PBS (1mg/ml, 100 μl), vortexed and sonicated until fully dissolved. 10 μl aliquot was taken at this stage as input. Then probe-labelled proteins were pulled-down using avidin-coated magnetic beads. After equilibrating the beads in 0.2% SDS/PBS, 10 μl of bead solution was added to samples, vortexed and incubated for 2-3 hours at room temperature. After brief centrifugation, samples were placed on magnet for 1 min and supernatant was removed carefully. 10 μl aliquot was taken at this stage as supernatant. Following several washes in 0.2% SDS/PBS, beads were boiled at 100°C for 5 min in the presence of Laemmli loading buffer containing β-mercaptoethanol and placed on magnet for 1 min. Probe-enriched samples were then transferred to a new tube and resolved by 12% gel SDS-PAGE. To check whether click reaction worked, TAMRA in-gel fluorescence was visualised using a Typhoon Biomolecular Imager (λ_exc_= 552 nm, λ_em_=570 nm; Amersham GE Healthcare). Thus, proteins were transferred to PDVF membranes using an iBlot Gel Transfer (Life Technologies) for 1 hour at 100 V. Then membranes were blotted and developed as mentioned above (see *Western blot* section) using as primary antibodies anti-STAT3 (Cell Signalling, #124H6) and anti-GAPDH (Abcam, #Ab9485).

### Gene expression analysis of ALDH+/- populations in ER+ metastatic breast cancer

Whole transcriptomic datasets for ALDH+ and ALDH- populations from ER+ metastatic patient-derived samples were available here (30). Briefly, ER+ advanced breast cancer patients were drained of metastatic fluids (ascites or pleural effusions) due to discomfort. Breast cancer cells were isolated following the methodology described elsewhere (19) and then stained using the ALDEFLUOR assay. After FACS sorting, RNA was extracted and the whole transcriptome of ALDH+ and ALDH-populations was assessed using Affymetrix Whole-Transcript Human Gene 1.0 ST Array (Affymetrix, Thermo Fisher Scientific) (30). Four patient samples were selected on the basis of being ER+PR+HER2- and having received endocrine treatment prior to collection. Differentially expressed STAT3-related genes (upstream and downstream) between ALDH+ and ALDH-cells were identified using Ingenuity Pathway Analysis (Qiagen).

### Quantitative Real-Time PCR using the Biomark HD System (Fluidigm)

Six metastatic patient-derived samples were treated in adherence for 72h in the presence of 5 μM SFX-01 or water (vehicle control) in DMEM/F-12 GlutaMAX medium (GIBCO) containing 10% foetal bovine serum (FBS; GIBCO), 10 mg/ml insulin (Sigma-Aldrich), 10 mg/ml hydrocortisone (Sigma-Aldrich), and 5 ng/ml epidermal growth factor (EGF; Sigma-Aldrich). Total RNA was then extracted using RNeasy Mini Kit (Quiagen, #74104). RNA quality, integrity and concentration were assessed using the Qubit RNA HS assay and the Agilent Bioanalyzer. Gene expression was evaluated with the 48.48 IFC Dynamic Arrays (Fluidigm Corporation) using standard Taqman Assays as per protocol (PN 68000089 H1). Taqman Gene Expression Assays used: *CTTN* (Hs01124232_m1), *MUC1* (Hs00159357_m1), OSMR (Hs00384276_m1), PBX1 (Hs00231228_m1), *PGK1* (Hs99999906_m1), *SDHA* (Hs00188166_m1) (Thermo Fisher Scientific). cDNA was prepared with Reverse Transcription Master Mix (Fluidigm, #100-6297). Samples were incubated on a thermal cycler (MJ Research) for 5 min at 25°C, 30 min at 42°C and 5 min at 85°C. For the pre-amplification step (PN 100-5876 C2), a diluted pool of gene expression assays was prepared with the Dilution Reagent (10 mM Tris-HCl, pH 8.0, 1 mM EDTA, Fluidigm, #100-8726) and mixed with the Preamp Master Mix (Fluidigm, #100-5744) and cDNA. Samples were then incubated in a T100 thermal cycler (BioRad) for 2 min at 95°C followed by 14 cycles of 15 sec at 95°C and 4 min at 60°C. After cycling, resulting samples were diluted 1:5 with Dilution Reagent. Then each 20X Taqman Gene Expression Assay was diluted in 2X Assay Loading Reagent (Fluidigm, #100-7611). Sample pre-mix was made by combining Taqman Universal PCR Master Mix 2X (Life Technologies, #4304437), 20X GE Sample Loading Reagent (Fluidigm, #100-7610) and pre-amplified cDNA. Following hydraulic chip priming, the diluted assays and samples were transferred into the appropriate IFC inlets and loaded with the IFC MX Controller (Fluidigm). Then the loaded chip was placed in the Biomark HD instrument (Fluidigm) and run the GE 48×48 specific protocol, 96.5 °C for 10 min followed by 40 cycles at 96 °C for 15 sec and 60 °C for 1 min. The data were normalised to average of housekeeping genes and ΔCt, ΔΔCt and fold change values were calculated for each metastatic sample.

### Statistical analysis

Unless otherwise stated, statistical analyses were carried out by a two-tailed Student’s t-test. A p-value≤0.05 was considered to be statistically significant. Error bars represent the standard error of mean (SEM) of at least three independent experiments. Data is shown as mean ± SEM.

Survival analysis was performed on patient datasets using either the KMplotter online tool (23) or performing Cox proportional hazards tests for all possible points-of-separation using the survivALL R package (31).

## ACKNOWLEDGEMENTS

We are grateful to Breast Cancer Now (MAN-Q2) for funding this research. B.M.S., R.B.C. and S.J.H. are supported by the NIHR Manchester Biomedical Research Centre (IS-BRC-1215-20007). Evgen Pharma and The Christie Charitable Fund also supported part of the study.

## SUPPLEMENTARY FIGURE LEGENDS

**TABLE S1:**
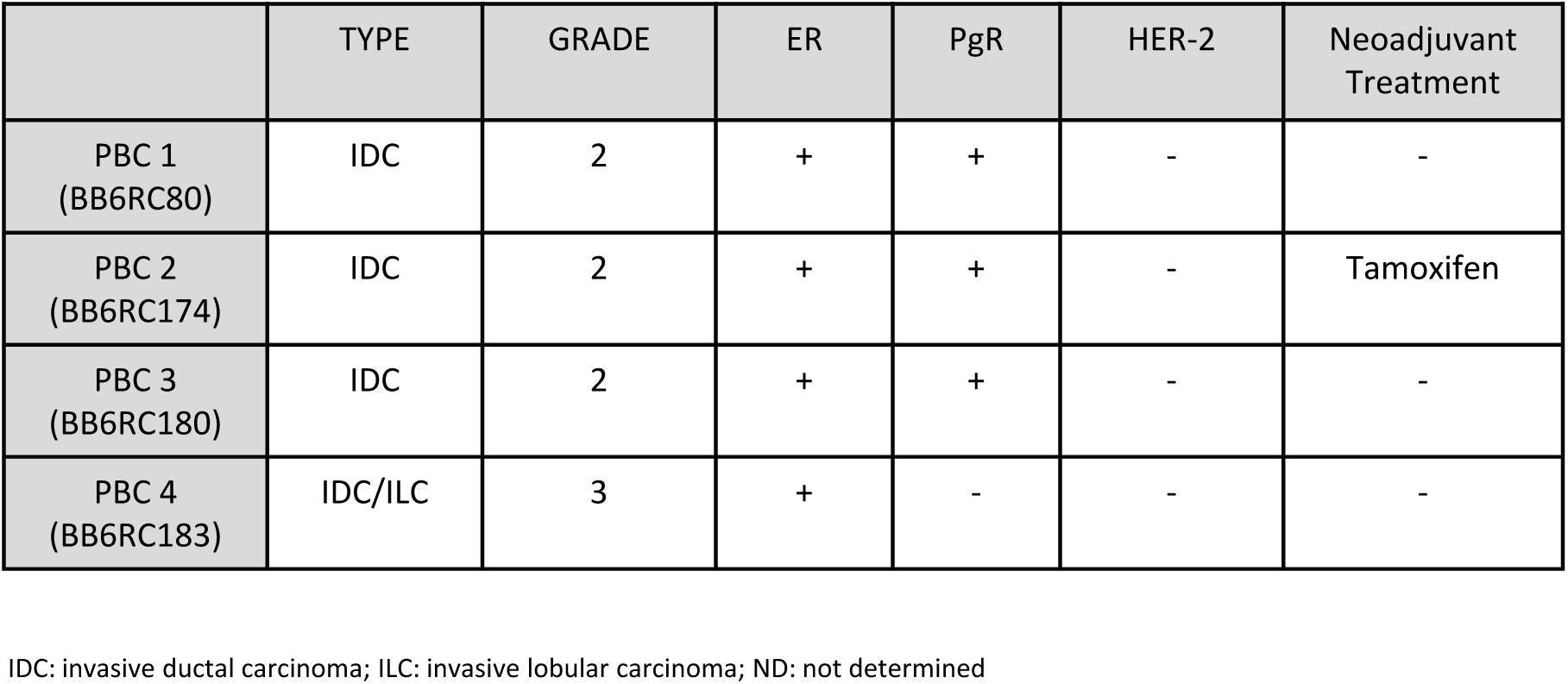
Characteristics of ‘early’ patient-derived tumours used in the study. PBC – primary breast cancer.

**TABLE S2:** Characteristics of ‘late’ metastatic endocrine therapy-treated patient-derived tumours used in the study. Met BC – metastatic breast cancer.

|  | TYPE | GRADE | ER | PgR | HER-2 | Chemotherapy | Hormonal therapy | Targeted therapy |
| --- | --- | --- | --- | --- | --- | --- | --- | --- |
| Met BC 1<br>(BB3RC44)<br>Met BC 2<br>(BB3RC59)<br>Met BC 3<br>(BB3RC61)<br>Met BC 4<br>(BB3RC66) | ILC | 2 | + | + | - | EOX<br>Capecitabine<br>Paclitaxel | Letrozole<br>Tamoxifen<br>Fulvestrant<br>Exemestane<br>Diethylstilbestrone | Zoledronic<br>acid/pamidronate |
| Met BC 5<br>(BB3RC60) | ILC | 2 | + | + | - |  | Fulvestrant<br>Anastrozole |  |
| Met BC 6<br>(BB3RC68) | IDC | 2 | + | N/K | - | FEC<br>Capecitabine<br>Paclitaxel | Tamoxifen<br>Anastrozole<br>Fulvestrant |  |
| Met BC 7<br>(BB3RC69)<br>Met BC 8<br>(BB3RC70) | ILC | 2 | + | + | - | ECF | Tamoxifen<br>Letrozole<br>Anastrozole<br>Fulvestrant | Zoledronic<br>acid/pamidronate |
| Met BC 9<br>(BB3RC72) | N/K | N/K | + | + | - | Lapatinib<br>Capecitabine<br>Paclitaxel<br>EC<br>Eribulin | Letrozole<br>Fulvestrant<br>Anastrozole<br>Exemestane<br>Tamoxifen | Zoledronic acid<br>Everolimus |
| Met BC 10<br>(BB3RC74) | ILC | 2 | + | + | - |  | Anastrozole<br>Tamoxifen |  |
| Met BC 11<br>(BB3RC77)<br>Met BC 12<br>(BB3RC77b) | ILC | 2 | + | + | - | Paclitaxel<br>Capecitabine<br>Eribulin | Letrozole<br>Exemestane<br>Fulvestrant | Ibandronate |
| Met BC 13<br>(BB3RC81) | IDC | 2 | + | + | - | FEC<br>Paclitaxel<br>Capecitabine | Tamoxifen<br>Anastrozole |  |
| Met BC 14<br>(BB3RC82) | IDC | 2 | + | + | - | TAC<br>Capecitabine<br>Paclitaxel | Tamoxifen<br>Anastrozole<br>Letrozole<br>Goserelin | Ibandronate<br>Zoledronic acid |
| Met BC 15<br>(BB3RC83) | IDC | 3 | + | + | - | FEC<br>Docetaxel<br>Vinorelbine<br>Eribulin | Tamoxifen<br>Anastrozole<br>Fulvestrant | Zoledronic acid |
| Met BC 16<br>(BB3RC89) | IDC | N/K | + | N/K | - | EC<br>Taxol<br>Capecitabine<br>Eribulin | Exemestane | Everolimus |
| Met BC 17<br>(BB3RC91)<br>Met BC 18<br>(BB3RC91A) | ILC | 2 | + | + | - | FEC<br>Docetaxel | Anastrozole<br>Letrozole<br>Fulvestrant<br>Exemestane | Zoledronic acid<br>Everolimus |
| Met BC 19<br>(BB7023) | IDC | N/K | + | + | - | Capecitabine<br>Paclitaxel<br>Epirubicin<br>Vinorelbine<br>Eribulin | Goserelin<br>Anastrozole<br>Fulvestrant<br>Exemestane |  |
| Met BC 20<br>(BB7103) | ILC | 2 | + | - | - | FEC<br>Paclitaxel<br>Capecitabine<br>Eribulin | Letrozole<br>Fulvestrant | Denosumab |
| Met BC 21<br>(BB7121) | ILC | 2 | + | + | - | E-CMF<br>Paclitaxel | Tamoxifen<br>Fulvestrant | AZD5363<br>SFX-01 (STEM trial) |

IDC: invasive ductal carcinoma; ILC: invasive lobular carcinoma; 5-FU: 5-Fluorouracil; EC: Epirubicin, Cyclophosphamide; FEC: 5-FU, Epirubicin, Cyclophosphamide; FEC-T: 5-FU, Epirubicin, Cyclophosphamide, Docetaxel; CMF: Cyclophosphamide, Methotrexate, 5-FU; TAC: Docetaxel, Doxorubicin, Cyclophosphamide; ECF: Epirubicin, Cisplatin, 5-FU; EOX: Epirubicin, Oxaliplatin, Capecitabine; E-CMF: Epirubicin-Cyclophosphamide, Methotrexate, 5-FU; N/K: not known
NB: ER, PgR and HER-2 were assessed in the primary breast cancer tissue sample.

**Supplementary figure 1.**
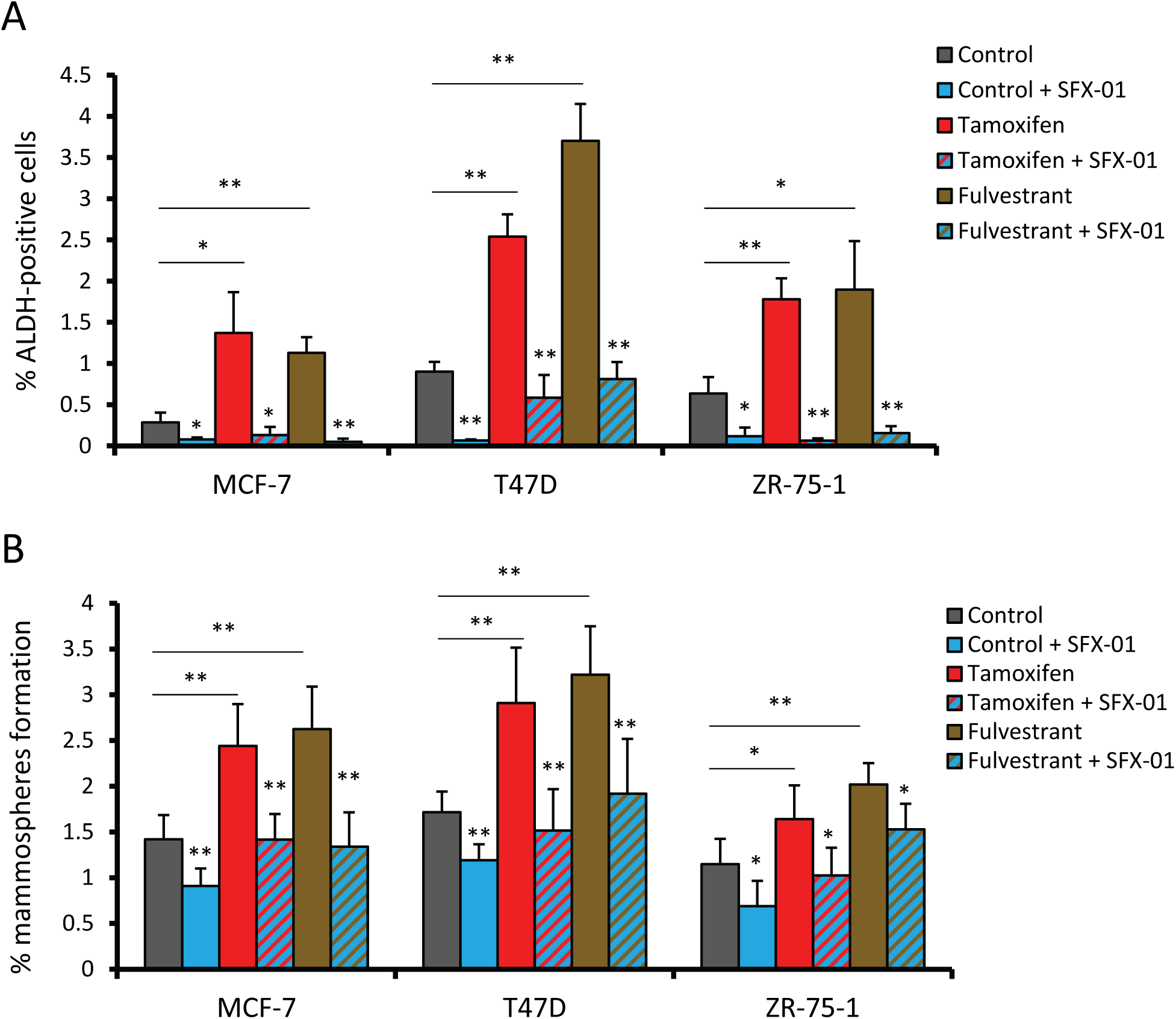
A) Mammosphere formation efficiency of MCF-7, T47D and ZR75-1 cells. Cells were pre-treated in adherence with ethanol (Control), tamoxifen (1 μM) or Fulvestrant (0.1 μM) in the presence of SFX-01 (5 μM) or water for 72 hours, and then plated at low density (200 cells/cm^2^) into 6-well plates and allowed to grow in suspension for 5 days. Mammosphere formation efficiency was calculated by dividing the number of mammospheres formed (≥ 50μm diameter) by the original number of single cells seeded and is expressed as the mean percentage of mammosphere formation ± standard deviation. B) FACS analysis of ALDH1 enzymatic activity assessed by the ALDEFLUOR assay in MCF-7, T47D and ZR75-1 cells. Graphs show the mean percentage of ALDH-positive cells (± standard deviation) for each cell line grown and treated as in A). After 72 hours of treatment, ALDEFLUOR assay was performed. p values refer to SFX-01 treatment compared to respective non-SFX-01 bars, * p < 0.05, ** p < 0.01

**Supplementary figure 2.**
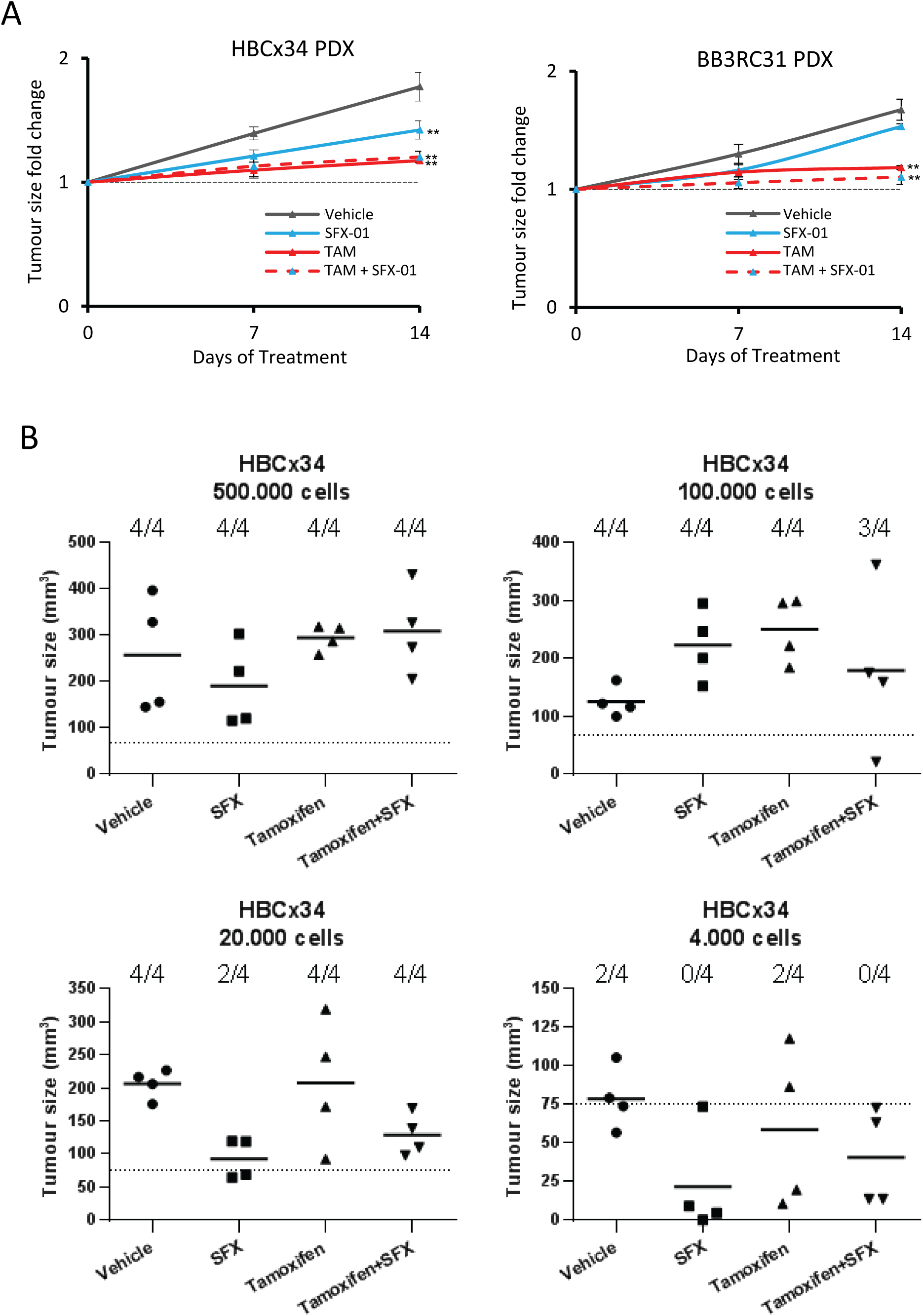
A) Early (HBCx34) and metastatic (BB3RC31) PDXs tumor size variation over 14 days *in vivo* treatments with SFX-01 (300mg/kg/day, oral gavage) in the presence or absence of tamoxifen (10mg/kg/day, oral gavage). Tumor size was determined every 3-4 days and averaged for each week. Fold change was calculated by dividing the tumor size by the size of the respective tumor at day 0. B) Graphs representing tumor size at day 90 after cell injection for each cell number tested. HBCx34 cells were pre-treated *in vivo* for 14 days. Experiments (N=4 per condition) were carried out in NSG mice injected subcutaneously with 500,000, 100,000, 20,000 and 4,000 cells. 90-day slow release estrogen pellets were implanted subcutaneously into mice two days before cell injection (0.72 mg, Innovative Research of America). Positive tumor growth was assessed by determining the mice bearing a tumor greater than 75 mm^3^ and is represented as mice positive for growth/mice tested. ** p < 0.01

**Supplementary figure 3.**
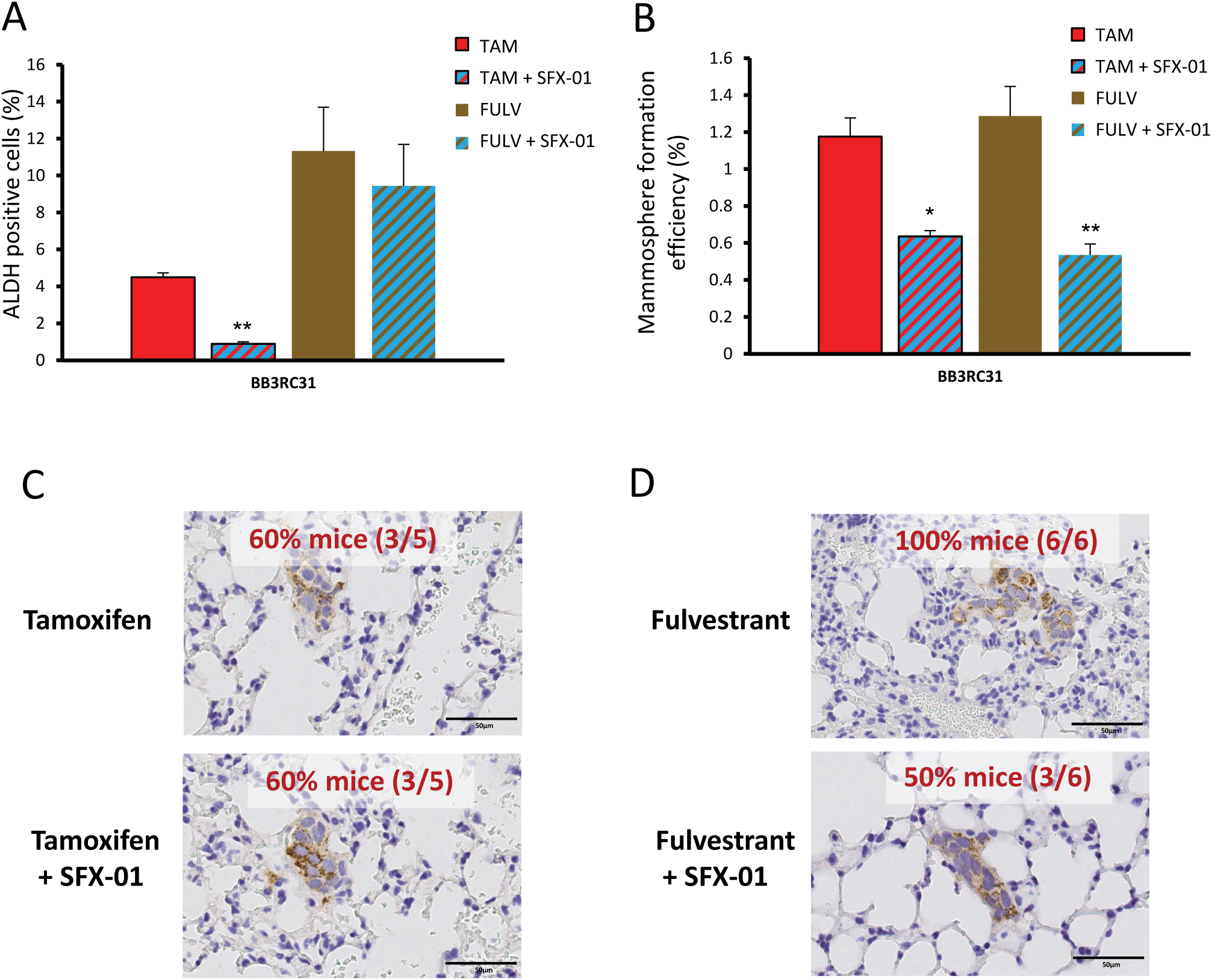
BB3RC31 PDX treated *in vivo* for 56 days with tamoxifen (10mg/kg/day, oral gavage) or fulvestrant (200mg/kg/week, subcutaneous injection) in the presence or absence of SFX-01 (300mg/kg/day, oral gavage). A) Percentage of ALDH-positive cells was determined with ALDEFLUOR assay. ALDH-positive cells were discriminated from ALDH-negative cells using the ALDH inhibitor, DEAB. B) Mammosphere formation efficiency was determined on day 7-9 and calculated by dividing the number of mammospheres formed (≥ 50μm diameter) by the original number of single cells seeded (500 cells/cm^2^) and is expressed as the mean percentage of mammosphere formation. C-D) Mice lungs were stained with anti-human mitochondrial antibody and micrometastases with at least 10 cells were counted. Percentage of mice bearing micrometastases for each tamoxifen (C) and fulvestrant (D) treatment group is shown. Data is represented as mean ± SEM. * p < 0.05; ** p < 0.01

**Supplementary figure 4.**
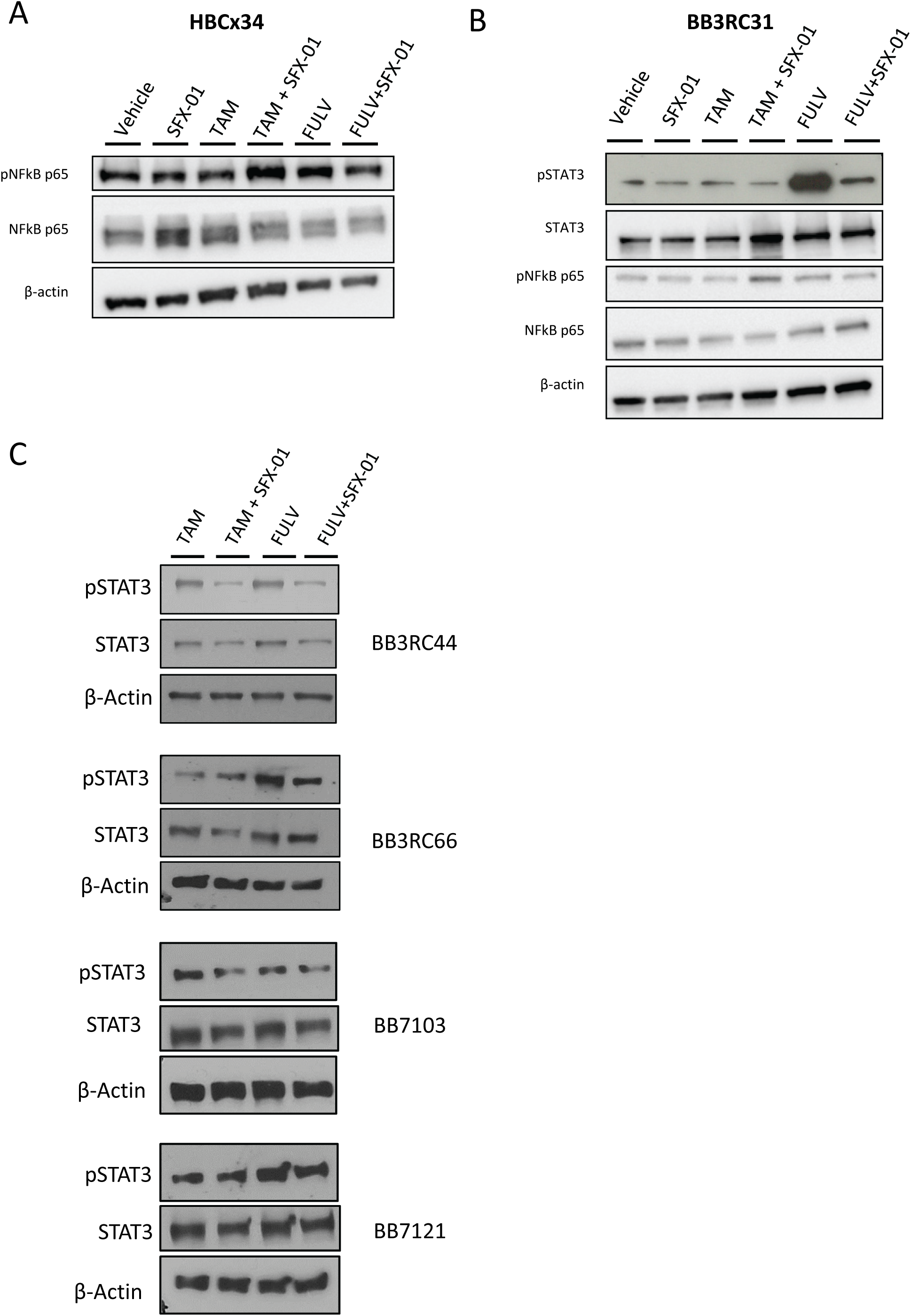
A) phospho-NFkB p65 and total NFkB p65 protein expression levels determined by Western Blot in HBCx34 PDX treated *in vivo* for 56 days with tamoxifen or fulvestrant in the presence or absence of SFX-01. β-actin was used as a reference for the loading control. B) phospho-STAT3, total STAT3, phospho-NFkB p65 and total NFkB p65 protein expression levels determined by Western Blot in BB3RC31 PDX treated *in vivo* for 56 days with tamoxifen or fulvestrant in the presence or absence of SFX-01. C) phospho-STAT3 and total STAT3 protein expression levels in four metastatic patient-derived samples treated for 72 hours with tamoxifen or fulvestrant in the presence or absence of SFX-01.

**Supplementary figure 5.**
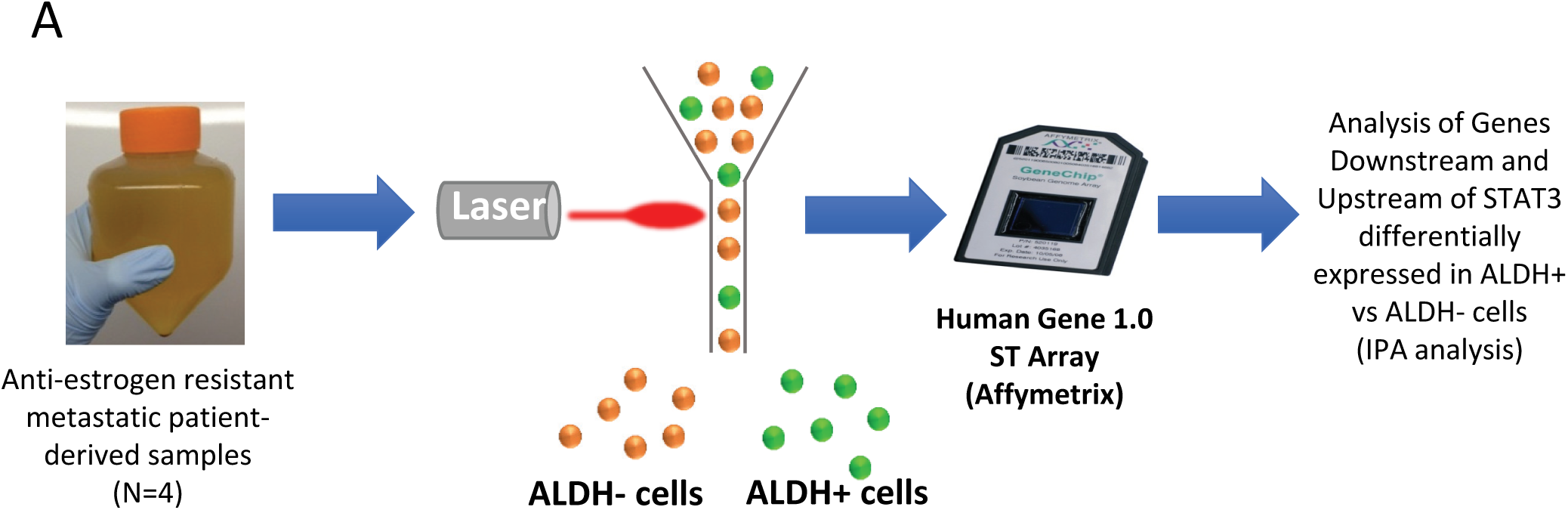
A) Schematic overview of the experimental approach used to profile gene expression of ALDH+ and ALDH- cells from metastatic patient samples. Ingenuity Pathway Analysis (IPA) software was employed to determine STAT3-related genes differentially expressed in ALDH+ cells.

